# Physiologically refined cell culture conditions uncover oncogene-dependent metabolic signatures in Ewing sarcoma spheroids

**DOI:** 10.1101/2024.07.05.602211

**Authors:** A. Katharina Ceranski, Martha J. Carreño-Gonzalez, Anna C. Ehlers, Zuzanna Kolodynska, Endrit Vinca, Tobias Faehling, Kimberley M. Hanssen, Philipp Poeller, Shunya Ohmura, Florencia Cidre-Aranaz, Almut Schulze, Thomas G. P. Grünewald

## Abstract

Ewing sarcoma (EwS) cell line culture largely relies on standard techniques, which do not recapitulate physiological conditions. Here, we report on a physiologically improved, feasible, and cost-efficient EwS cell culture technique employing an advanced medium composition, reduced fetal calf serum, and spheroidal growth. Functional *in vitro* assays and transcriptome profiling demonstrated that these refined conditions better recapitulate proliferation rates of patient tumors as well as hypoxic conditions relevant for EwS pathophysiology. Moreover, transcriptional signatures associated with the oncogenic activity of the EwS-specific FET::ETS fusion transcription factors in the refined culture conditions were shifted from proliferative towards metabolic gene signatures. The herein presented optimized physiological EwS cell culture technique provides a broadly applicable approach for enhanced *in vitro* modeling relevant to advancing EwS research and the validity of experimental results.

**MOTIVATION:** Cell culture remains the main platform to model EwS for research purposes. Yet, concerns exist about the limitations of standard *in vitro* techniques to adequately reflect physiological conditions. In this study, we refined EwS cell culture methods to increase modeling capacity while ensuring a practical and cost-effective handling, thereby broadening their applicability within the scientific community.

## INTRODUCTION

Ewing Sarcoma (EwS) is an aggressive pediatric bone and soft tissue cancer that is genetically characterized by *FET::ETS* fusion oncogenes encoding aberrant transcription factors, mostly *EWSR1::FLI1*^1^. The clinical outcome of many patients, especially that of patients with metastatic disease at diagnosis, remains limited (typically <30% overall survival rates)^1^. There is consensus in the scientific community that physiological conditions are required to improve the fidelity of preclinical disease models, which may in turn lead to the identification of more effective therapies^2–4^. In fact, the main preclinic EwS tumor model involves the culture of EwS cell lines in standard techniques, employing a two-dimensional (2D) monolayer set-up and traditional cell culture media^5^. However, 2D monolayer culture does not represent architectural and mechanical aspects of the EwS physiological tumor niche and has been identified to limit the translatability of preclinical research advances into improved therapies for EwS patients^6,7^. Consequently, various three-dimensional (3D) cell culture techniques, advanced scaffold- modeling of the bone extracellular matrix (ECM), flow perfusion techniques, and microfluidic systems have been introduced to improve EwS model accuracy^2–4,7^. Other approaches for more accurate preclinic EwS models are the genetic engineering of EwS cells from mesenchymal stem cells^8^, the use of various types of xenografted animal models^9,10^ and the attempt to create genetically engineered mouse models of EwS, which has not been successful until today^11^. Yet, from a practical standpoint, the complexity of any experimental technique will largely determine their experimental throughput, cost-effectiveness, broad applicability, and efforts needed for data acquisition^3,12^. Besides the improvements addressing the architectural aspects of the standard EwS cell culture technique, evidence suggests that physiologically refined cell culture media like human plasma-like medium (HPLM)^13^ and Plasmax^14^ are essential to enhance the fidelity of *in vitro* cell culture techniques^5,15,16^. Yet, physiological cell culture media^13,14^ have not been implemented for the culture of EwS cell lines until to date. Furthermore, although it is well-known that additives such as fetal calf serum (FCS) introduce nutrients, growth factors, and hormones at unknown concentrations to the culture medium^5,17,18^, they are typically applied in high and empiric concentrations, which can compromise reproducibility of experiments. In fact, little is known on whether EwS cell culture could also be feasible at reduced FCS concentrations that would better comply to the 3R principles (reduction, refinement, replacement) thereby contributing to the promotion of animal-welfare in the experimental space^17,18^.

In this paper, we present a physiologically refined, simple, and inexpensive technique for EwS cell culture amenable to large-scale use. We chose a spheroid approach^19^ and applied an advanced medium based on the formulations of HPLM^13^ and Plasmax^14^ which mimic nutrient concentrations observed in the human plasma, as well as carefully adapted reduced FCS concentrations to increase modeling relevancy of EwS cell culture. Using functional *in vitro* assays and transcriptome profiling, we show that these improved physiological cell culture conditions better mimic proliferation rates of patient tumors as well as hypoxic conditions that shape the EwS tumor microenvironment^20,21^. Moreover, we demonstrate that the refined culture techniques have a pervasive effect on the baseline transcriptomes of EwS cells, and significantly impact the transcriptional outputs of EWSR1::ETS fusion oncogenes.

## RESULTS

### Establishing physiologically optimized cell culture conditions for EwS cell lines

To improve the physiological relevancy of the EwS standard culture conditions, we addressed the following three technical aspects in the refined EwS cell culture technique: medium composition, FCS concentration, and growth structure (**Figure 1A**). In the first step, two established physiological medium compositions, HPLM^13^ and Plasmax^14^, were combined for a newly formulated physiological medium, hereafter called HPLMax (**Figure 1B**, **Supplementary Table S1**, see Methods). EwS cell lines were considered adapted after four weeks of culturing in HPLMax and were then characterized phenotypically. As a second step, to evaluate the minimal necessary FCS concentration that did not compromise EwS cell viability, we performed a resazurin proliferation assay for five selected EwS cell lines cultured in Roswell Park Memorial Institute 1640 Medium (RPMI) medium with FCS concentrations ranging from 10% (standard) to 1%. The human EwS cell lines A-673, MHH-ES-1, SK-N-MC, TC-71 (all *EWSR1::FLI1*-positive) and TC-106 (*EWSR1::ERG*-positive) were selected for study since they are widely used in EwS research and as they reflect the two major *EWSR1::ETS* fusions found in ∼95% of patients^1^. Of note, all cell lines were previously engineered to harbor doxycycline (dox)-inducible shRNAs to target the respective *FET::ETS* fusions^22^. Interestingly, FCS supplementation could be reduced to 7% in the five EwS cell lines without significantly decreasing their proliferation (**Figure 1C, Supplementary Figure S1**). In fact, 7% FCS supplementation resulted in even slightly accelerated proliferation for some cell lines (SK-N- MC/TR/shEF1 and TC-71/TR/shEF1, see also **Supplementary Figure S1**). The 7% FCS optimized condition was further validated by manual cell counting using standardized hemocytometers and Trypan-blue exclusion methods in both RPMI and in the physiological HPLMax medium (**Figure 1D**). Moreover, manual cell counting demonstrated no notable change in EwS cell viability at 7% FCS in both media (**Figure 1E**). We concluded that 7% FCS supplementation is optimal for these EwS cell lines to provide the required growth factors while allowing a 30% reduction of this animal product required for experimentation and its entailed drawbacks^5,17,18^. While proliferation rates of all EwS cell lines remained stable within both media (RPMI and HPLMax) despite the 30% decrease in FCS supplementation (**Figure 1D**), we did observe a slower proliferation in the five EwS cell lines when comparing the doubling time in HPLMax to RPMI under an equal medium exchange frequency (**Figure 1F**). This result is consistent with the fact that conventional media were designed to maximize the growth rates of cultured cells^13,14^ and was also observed for proliferation of hematological and epithelial cancer cell lines in HPLM^13,23^. Concerning the viability of the five EwS cell lines, similar dead cell counts were found in RPMI and HPLMax (**Supplementary Figure S1**). Regarding the EwS cell line morphology, no striking changes were observed *in vitro* between the culture in RPMI and HPLMax (**Figure 1G, Supplementary Figure S1**). As a final step in the optimization, the traditional 2D monolayer culture pattern was replaced with a spheroid approach (**Figure 1H**) by adjusting a published protocol for agar-coated low attachment spheroid culture^19^ (see Methods). This approach allows for better mimicking of the tumor architecture (**Figure 1H**), cell-cell interactions, and spatial gradients of oxygen and nutrients, which are all essential components of the tumor microenvironment^3^.

**Figure 1:**
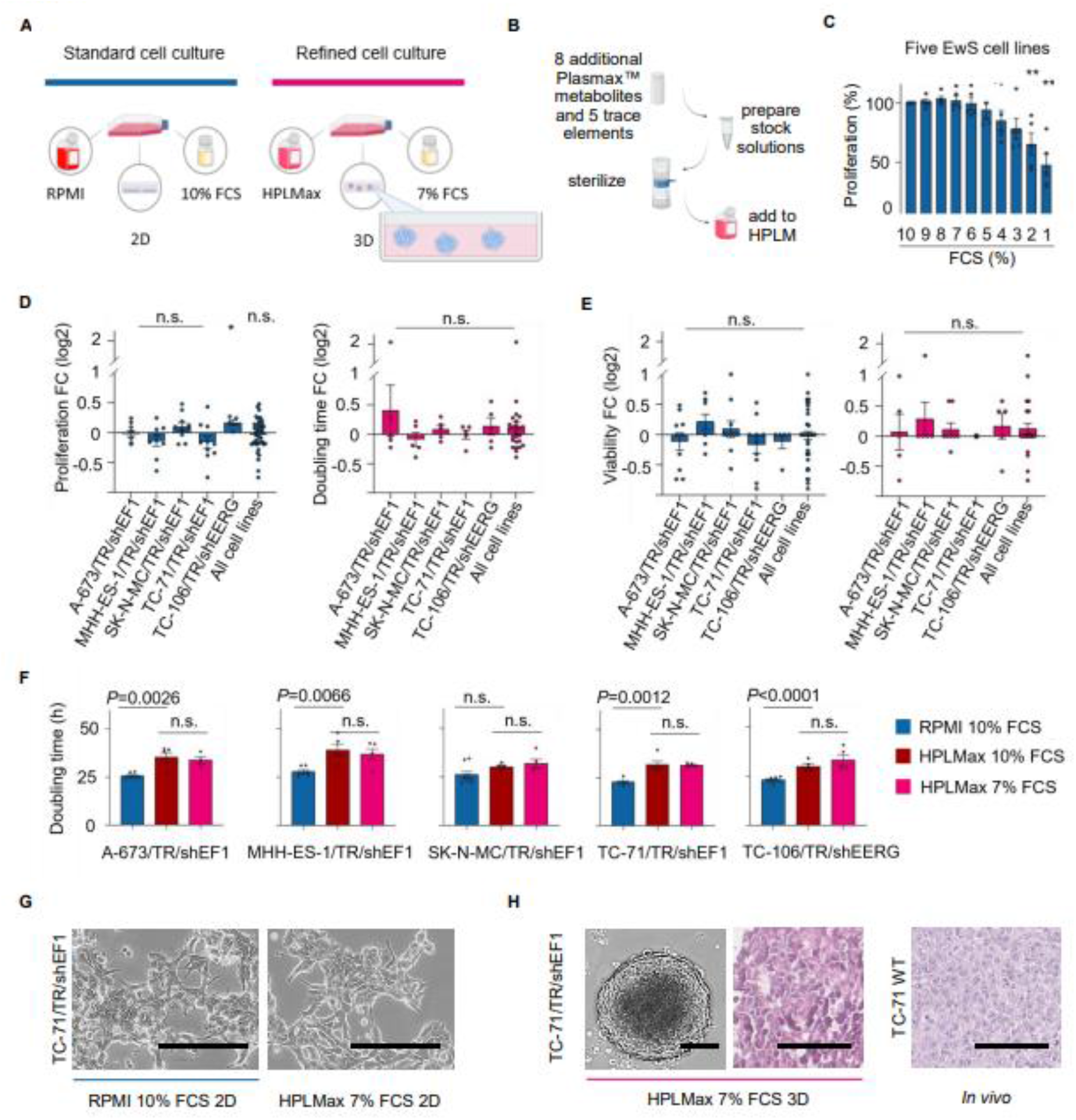
Establishing physiologically optimized cell culture conditions for EwS cell lines **A** Overview on EwS standard cell culture technique and the physiologically refined technique. **B** Workflow to create HPLMax (see also **Table S1**). **C** Proliferation assessment by resazurin cell viability assays in EwS cell lines in RPMI with FCS supplementation from 10% to 1%. Values were normalized to the proliferation in 10% FCS. Pooled data of five representative cell lines with each dot representing n≥5 biologically independent replicates of one cell line. Vertical bars represent means and whiskers the standard error of the mean (SEM). Two-sided Mann-Whitney test. \*\**P*<0.01, \*\*\**P*<0.001 (see also **Figure S1**). **D** Viable cell counts using Trypan blue dye exclusion method in 7% FCS depicted as fold change to those in 10% FCS (log2) in RPMI (left) and HPLMax (right). Each dot represents a single biologically independent experiment. Vertical bars represent means and whiskers the SEM. Two-sided one sample t-test. \**P* < 0.05, n.s. = non-significant. **E** Dead cell counts using Trypan blue dye exclusion method in 7% FCS depicted as fold change to those in 10% FCS (log2) in RPMI (left) and HPLMax (right). Each dot represents a single biologically independent experiment. Vertical bars represent means and whiskers the SEM. Two-sided one sample t-test. **F** Doubling time of the five selected EwS cell lines in RPMI, 10% FCS (left bar), HPLMax, 10% FCS (middle bar) and HPLMax, 7% FCS (right bar) by trypan blue counting of viable cells. Each dot represents a single biologically independent experiment. Vertical bars represent means and whiskers the SEM. Two-sided unpaired t-test (see also **Figure S1**). **G** Brightfield microscopy images of TC-71/TR/shEF1 cell lines grown in RPMI and 10% FCS (left) and in HPLMax and 7% FCS (right). Scale bars = 200 μm, 10× magnification (see also **Figure S1**). **H** TC- 71/TR/shEF1 spheroids grown in agar-coated T25 flasks with HPLMax and 7% FCS for 8 d, light microscopy image (left), scale bar = 125 μm, 4× magnification. Hematoxylin/Eosin (H&E) staining of TC-71/TR/shEF1 spheroids grown as described above and treated with doxycycline for 96 h for KD of the fusion oncogene (middle), scale bar = 100 μm, 40× magnification. H&E staining of mice *in vivo* xenografts of TC-71 wildtype cells (right), scale bar = 100 μm, 40× magnification.

### Physiologically refined EwS cell culture condition preserves cell line characteristics but alters transcriptional programs related to proliferation, differentiation, and metabolism

To further characterize the EwS cells in the refined culture technique and to validate this set- up, we performed DNA microarray gene expression analyses in four EwS cell lines (MHH-ES- 1/TR/shEF1, SK-N-MC/TR/shEF1, TC-71/TR/shEF1, TC-106/TR/shEERG) that enable a dox- inducible and shRNA-mediated knockdown of either EWSR1::FLI1 or EWSR1::ERG, respectively^22^. An experimental overview is given in **Figure 2A**. Briefly, the Ewing Sarcoma Cell Line Atlas (ESCLA) transcriptome experiment from our laboratory in standard cell culture^22^ was recapitulated in the refined cell culture set-up with/without addition of dox for 96 h. Gene expression data were generated on the same platform (human Affymetrix Clariom D arrays), jointly preprocessed, and normalized with the former data^22^ to generate a harmonized dataset for direct comparison of refined and standard culture conditions. As shown in **Figure 2B** and **Supplementary Figure S2**, exploratory data analysis with all samples using principal component analysis (PCA) revealed that the captured variation was mainly driven by three aspects: the employed cell culture condition (PC1), the specific characteristics of the individual cell lines (PC2 – PC4), and the dox-dependent *EWSR1::ETS* expression levels (PC5). This illustrates that the applied cell culture technique fundamentally affects transcriptional programs of EwS cell lines, yet their principal characteristics are preserved in the refined set-up. Next, to further explore the effects of the cell culture conditions on baseline RNA transcription, we focused on culture conditions without *EWSR1::ETS* knockdown (dox–). Pre-ranked fast Gene Set Enrichment Analysis (fGSEA) with differentially expressed genes (DEGs) between the refined and standard culture condition followed by Weighted Gene Correlation Network Analysis (WGCNA) uncovered reduced cell-cycle- and proliferation-associated signatures as well as enrichment of ribosomal, translational, and spliceosomal processes in the physiologically refined culture conditions (**Figure 2C** and **2D**, **Supplementary Table S2**). Furthermore, we observed a strong dysregulation of canonical hallmark signatures as well as generic gene signatures involved in the regulation of differentiation (epithelial-to-mesenchymal transition, EMT), hypoxia, and metabolism in the refined set-up (**Figure 2D**). These data are in keeping with the results from the functional *in vitro* proliferation assays (**Figure 1G**) and the oxygen gradients that are a key feature of spheroid cultures^3^. Collectively, these results indicate that the physiologically refined culture technique preserves typical characteristics of EwS cell lines but impacts transcriptional programs related to proliferation, differentiation, metabolism and hypoxia.

**Figure 2:**
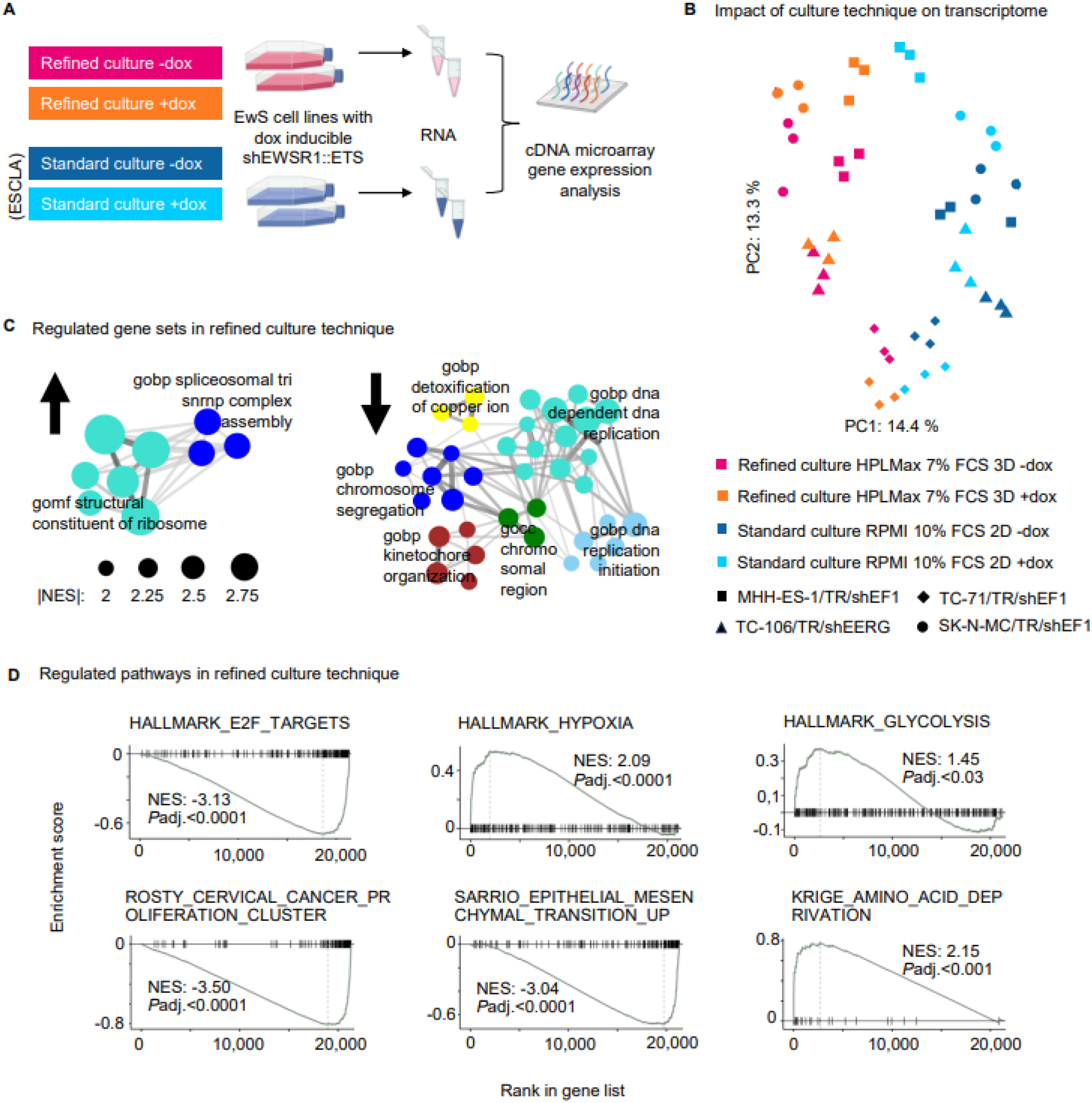
Physiologically refined EwS cell culture condition preserves cell line characteristics but alters transcriptional programs related to proliferation, differentiation, and metabolism **A** Experimental workflow to generate transcriptome data sets for EwS cell lines in physiologically refined cell culture conditions (pink and orange). The same experiment was previously performed for the Ewing Sarcoma Cell Line Atlas (ESCLA) in standard culture conditions by Orth *et al.*^22^ (blue and light blue). Data were analyzed via joint processing including background correction, normalization, and summarization (see Methods). **B** Principal Component Analysis (PCA) of the normalized data from (A) revealing captured variation driven by culture condition (PC1) and cell line characteristics (PC2). For captured variation by PC3 – PC5 see **Figure S2**. **C** Weighted correlation network analysis (WGCNA) based on pre- ranked fast gene set enrichment analysis (fGSEA) of differentially expressed genes (DEGs) from (A) using the average rank of expression fold change (FC). Displayed network analyses were based on significant gene sets with a maximum false discovery rate (FDR) |FDR| of 0.05 and a minimal normalized enrichment score (NES) |NES| of 2. See also **Table S2**. **D** Enrichment plots from fGSEA of DEGs from (A). Padj = Bonferroni adjusted *P*-value. C2 and Hallmark Gene Sets from the Human MSigDB Collections. See also **Table S2**.

### Physiologically refined cell culture conditions shape the transcriptional output of EWSR1::ETS

Since oncogenic EWSR1::ETS transcription factors are the most prominent genetic characteristic of EwS^1^, we subsequently investigated their transcriptional activity in the refined cell culture conditions in comparison to standard culture conditions. Dox-induced and shRNA-mediated knockdown of the fusions for 96 h resulted in both cell culture conditions in a similar and strong downregulation of a well-established EWSR1::ETS-related gene signature (‘Kinsey_targets_of_EWSR1::FLI1-fusion_down’, NES < –3) (**Figure 3A**). We next compared differentially regulated genes under EWSR1::ETS knockdown in the two culture techniques for shared EWSR1::ETS positively correlated and EWSR1::ETS negatively correlated genes in both culture conditions as well as genes uniquely regulated in either culture technique. As shown in **Figure 3B**, while there were notable overlaps in EWSR1::ETS regulated genes between the two conditions, there was also a large proportion (∼50%) of genes specific to each culture technique. Among the EWSR1::ETS regulated genes that were shared in both cell culture conditions, Gene Ontology (GO) analysis revealed that upregulated gene sets in EWSR1::ETS high conditions were predominantly associated with cell cycle progression (**Figure 3C, Supplementary Table S3**). Similarly, proliferation signatures were overrepresented among EWSR1::ETS correlated genes exclusively in standard cell culture condition (**Figure 3C, Supplementary Table S3**), which corresponds to previous reports that linked higher EWSR1::ETS expression in EwS to increased proliferation^24,25^. Most interestingly, genes that were EWSR1::ETS correlated exclusively in the refined culture condition were primarily categorized by GO analysis as participating in amino acid membrane transport (e.g. L-alanine and leucine import across plasma membrane, GO:1904273 and GO:0098713) (**Figure 3C**, **Supplementary Table S3**). Hence, this indicates EWSR1::ETS facilitates nutrient transport that is not apparent under supraphysiological concentrations of nutrients present in traditional cell culture media. For EWSR1::ETS negatively correlated genes, GO analysis retrieved differentiation processes that were reduced through upregulation of EWSR1::ETS in genes overlapping between the two media conditions as well as in genes unique to either condition (**Figure 3D, Supplementary Table S3**), thereby reflecting the known role of EWSR1::ETS in maintaining a dedifferentiated state^24^. Collectively, our data show that the activity of EWSR1::ETS fusions in EwS cell lines is shifted under more physiological cell culture conditions from a hyperproliferative state towards enhanced amino acid transport, whilst maintaining a de-differentiated state. As a reliance on extracellular nutrients under nutrient limiting conditions can be an exploitable vulnerability in cancer cells, these findings may have significant implications for future EwS research that aims for accurate discoveries of EWSR1::ETS regulated genes and drug targets.

**Figure 3:**
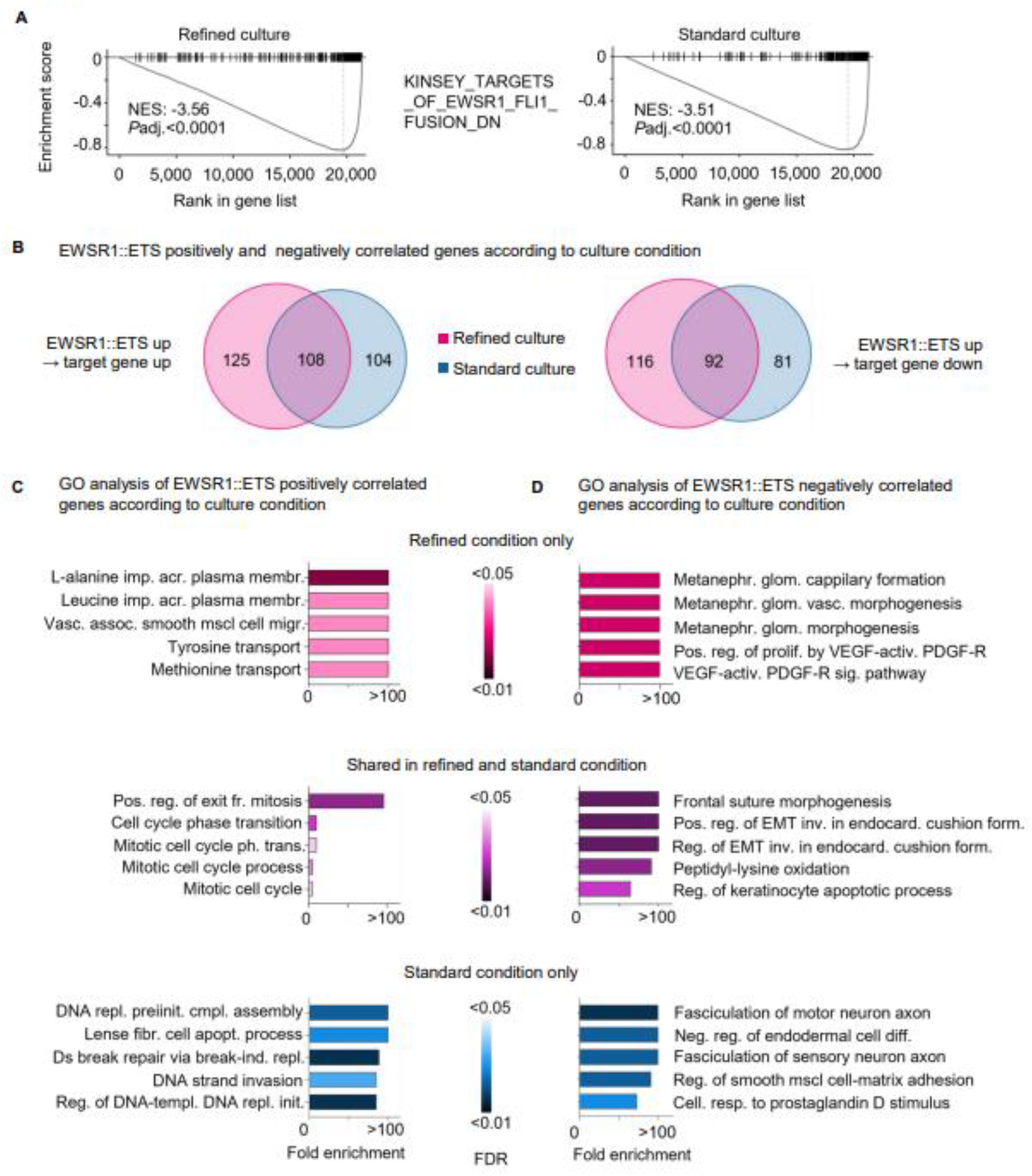
Physiologically refined cell culture conditions shape the transcriptional output of EWSR1::ETS **A** Enrichment plots from fGSEA of DEGs between EWSR1::ETS high vs EWSR1::ETS low expressing EwS cell lines in physiologically refined culture conditions (left) and standard culture conditions (right). C2 Gene Set from the Human MSigDB Collections. **B** Area- proportional Venn diagram showing overlap of DEGs, defined as having FDR adjusted *P*-value ≤ 0.05 and |log2 FC| ≥1.5, in refined culture conditions and standard culture conditions. Left side: EWSR1::ETS positively correlated DEGs, right side: EWSR1::ETS negatively correlated DEGs. See also **Table S3**. **C** Gene ontology (GO) analysis of DEGs from (B) showing top five enriched GO biological processes for each collection of DEGs: Upper row (pink) DEGs from refined culture conditions exclusively, middle row (violet) DEGs that are shared in both culture conditions, lower row (blue) DEGs from standard culture conditions only. Left column: EWSR1::ETS positively correlated DEGs, right column: EWSR1::ETS negatively correlated DEGs. See also **Table S3**.

### Cell culture conditions may influence expression of commonly used housekeeping genes

It is well-established that specific experimental conditions (e.g. cell culture techniques) can influence the expression levels of housekeeping genes (HKGs), which may introduce bias into studies based on qRT-PCR read-outs^26,27^. Indeed, when assessing our microarray data, we observed that *ribosomal protein lateral stalk subunit P0* (*RPLP0)*, a routinely used HKG in EwS research^22^, exhibited markedly different baseline expression levels between the standard and refined cell culture techniques. To nominate potential HKGs exhibiting minimal variation across cell culture conditions, we employed and adapted an approach described by de Jonge *et al.*^26^ (see Methods). As displayed in **Figure 4a**, the top three candidate HKGs were the *mitochondrially encoded cytochrome c oxidase I* (*MT-CO1*), *mitochondrially encoded NADH dehydrogenase 1* (*MT-ND1*), and *RNA component of 7SK nuclear ribonucleoprotein* (*RN7SK*), a small nuclear RNA (snRNA). Indeed, these three genes showed little variation as compared to *RPLP0* (**Figure 4b**). A comprehensive list of potential HKGs is provided in **Supplementary Table S4**. These data confirm the rationale that the choice of a suitable HKG must be tailored to the specific cell line and applied cell culture condition^26,27^.

**Figure 4:**
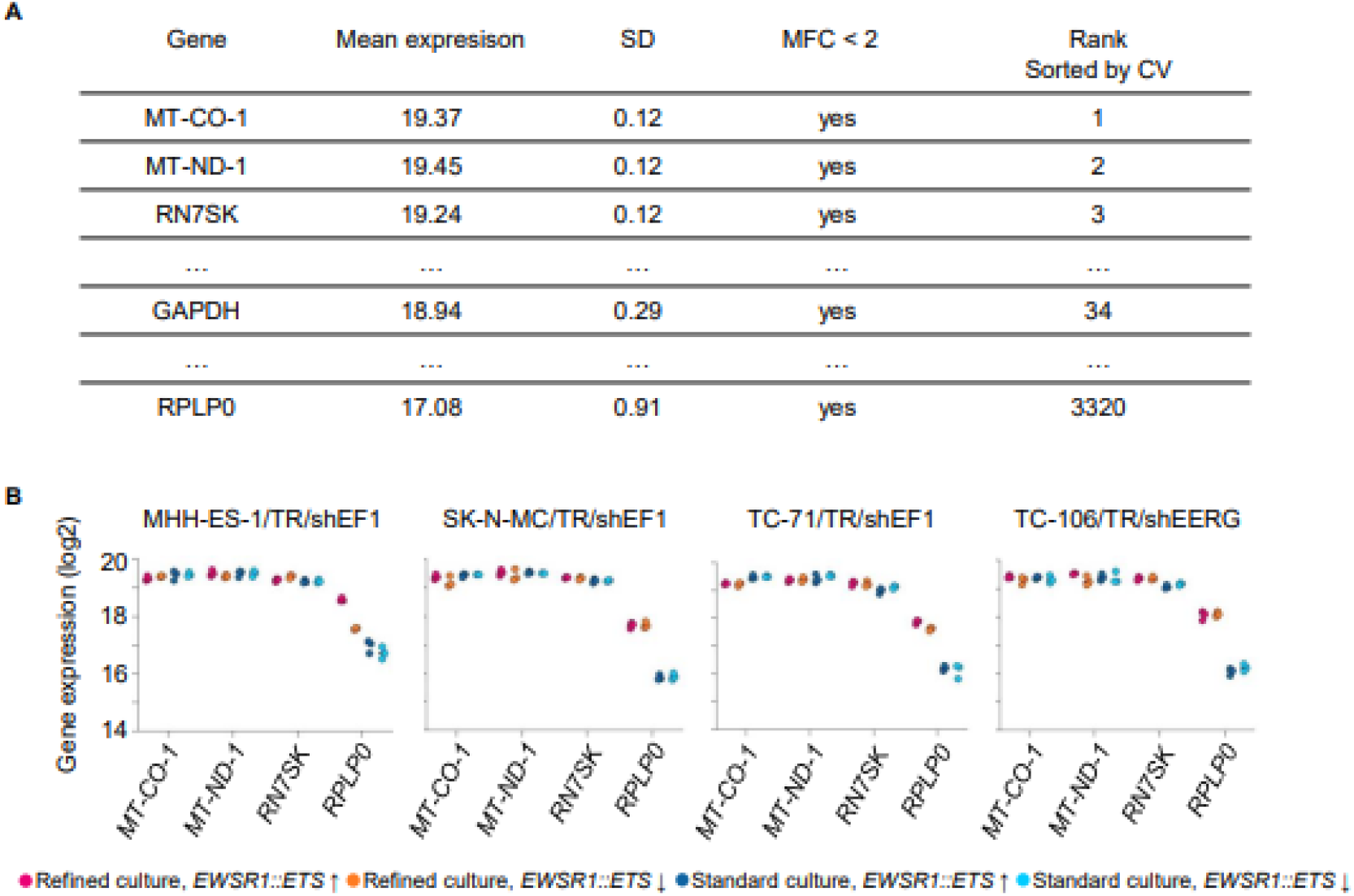
Cell culture conditions may influence expression of commonly used housekeeping genes **A** Table of the top three housekeeping gene (HKG) candidates and the routinely used HKGs *GAPDH* and *RPLP0*. HKG candidates were selected through the adapted approach from de Jonge *et al.*^26^ Mean expression value is defined as mean of the expression values of all samples from Fig. 2A including the four different conditions. SD = standard deviation, MFC = maximum FC, CV= coefficient of variation. See also **Table S4** and Methods. **B** Log2 transformed gene expression of HKG candidates from the microarray data in the four EwS cell lines from Fig. 2A.

## DISCUSSION

In this work, we pursue a practical approach to increase the physiological modeling relevancy of cell culture techniques for EwS cell lines. Our set-up was validated through a comparative transcriptome analysis derived from four EwS cell lines cultured in standard^22^ and physiologically refined culture conditions. Our analyses indicated that characteristic traits of the EwS cell lines, including EWSR1::ETS activity, were preserved in the physiologically refined set-up, while key aspects of EwS patient tumors, such as hypoxia^20,28^ and proliferation rates^29^, were better represented in the advanced culture technique. Hence, we consider the herein reported physiologically refined cell culture conditions as a suitable model for EwS *in vitro* studies. Most importantly, our results demonstrate that the employed cell culture conditions influence two aspects relevant for EwS research: i) transcriptional programs linked to a variety of cellular processes, such as proliferation, translation, differentiation, and metabolism, and ii) the activity of the EWSR1::ETS oncogenic transcription factors. Regarding aspect i), our data is consistent with previously published studies in that metabolism and proliferation are frequently altered when cells are cultured in physiological media^13,16,30^. This is also reflected in our data by the upregulation of ribosomal and translational processes when changing from standard to physiologically refined EwS cell culture conditions since translation and metabolism are tightly correlated both in normal and cancer cells^31,32^. Consequently, it is intriguing to further characterize EwS features with physiologically refined culture conditions by means of extended multi-omics profiling including the proteomic level.

Concerning aspect ii), our study provides insight into potential alterations of the metabolic landscape in EwS induced by activity of EWSR1::ETS oncoproteins in a physiologically relevant cell culture model. We hypothesize that observed altered expression of genes engaged in amino acid membrane transport in physiologic media is necessary to sustain proliferation of high EWSR1::ETS expressing EwS cell lines^24,25^ in physiological media. Interestingly, transcriptional regulation of genes involved in amino acid metabolic pathways in the context of proliferation in physiological media is in line with previous findings in other tumor entities^16,30^. Our data support the idea that altered metabolic pathways related to amino acid abundance are a characteristic trait of proliferating cells in physiological media. To fully characterize which amino acids and their metabolism are required for proliferation of EWSR1::ETS high expressing EwS cell lines, further studies on the interaction of FET::ETS oncogenes and the metabolism in EwS cells are needed. Finally, our data illustrate that the applied cell culture technique, as a part of the general experimental conditions, can influence the expression levels of routinely used HKGs^26,27^ – as was the case for *RPLP0*^22^ in this study.

In summary, our work underlines the necessity to improve the physiological fidelity of cell culture conditions, provides a pragmatic and cost-effective approach for the EwS field to overcome typical barriers for establishing these, and contributes a transcriptome data set generated in physiologically refined *in vitro* conditions as a valuable resource to the EwS research community.

## LIMITATIONS OF THE STUDY

The limitations of our refined cell culture technique for EwS cell lines involve primarily the natural restrictions of any spheroid-based technique. The ability of tumor cells to spontaneously form spheroids when cultured in low-attachment conditions is a prerequisite for this approach^3^. Further restrictions include the variability in size and diameter between individual spheres, if cultured on flat-bottom surfaces, the lack of a vasculature or dynamic flow system and, in our experimental set-up, the absence of ECM components^3^. On that account, alternative methods have been developed to mimic interactions of EwS cells with the ECM and mechanical forces^2–4,7^. These techniques could also be combined with the refined media composition and reduced FCS concentration that we propose in this report. Some of the limitations discussed above are a result of our intention to keep the cell culture technique simple, feasible and cost effective. It is due to this motivation that our physiologically refined cell culture method for EwS cell lines is amenable to routine and large-scale use and especially suitable for bulk analyses in a setting with increased physiological modeling relevancy.

## Supporting information

Ceranski_et_al_Supplementary_Table_S1

Ceranski_et_al_Supplementary_Table_S2

Ceranski_et_al_Supplementary_Table_S3

Ceranski_et_al_Supplementary_Table_S4

## ACKNOWLEDGMENTS

We thank Nadine Gmelin, Stefanie Kutschmann, and Felina Zahnow for excellent technical assistance. We thank Jia xiang Jin and Dr. Jing Li for fruitful discussion and advice during the project. We thank the Microarray Core Facility, German Cancer Research Center (DKFZ), for providing excellent Expression Profiling services. We thank Gaby Blaser from the DKFZ Light Microscopy Core Facility for valuable technical advice and gratefully acknowledge support by the DKFZ Light Microscopy Facility. The laboratory of T.G.P.G. is supported by grants from the Matthias-Lackas foundation, the Dr. Leopold und Carmen Ellinger foundation, the Deutsche Forschungsgemeinschaft (DFG 458891500), Dr. Rolf M. Schwiete foundation (2021- 007; 2022-031), the German Cancer Aid (DKH-7011411, DKH-70114278, DKH-70115315, DKH-70115914), the SMARCB1 association, the Ministry of Education and Research (BMBF; SMART-CARE and HEROES-AYA), the KiKa foundation, the Fight Kids Cancer foundation (FKC-NEWtargets), the KiTZ-Foundation in memory of Kirstin Diehl, the KiTZ-PMC twinning program, the German Cancer Consortium (DKTK, PRedictAHR), and the Barbara and Wilfried Mohr foundation. The laboratory of T.G.P.G. is co-funded by the European Union (ERC, CANCER-HARAKIRI, 101122595). Views and opinions expressed are however those of the authors only and do not necessarily reflect those of the European Union or the European Research Council. Neither the European Union nor the granting authority can be held responsible for them. A.C.E., T.F., and E.V. were supported by a scholarship of the German Cancer Aid and the German Academic Scholarship Foundation. E.V. and T.F. were additionally supported by a scholarship of the Heinrich F.C. Behr foundation, and E.V. by the Rudolf and Brigitte Zenner foundation. K.M.H. was supported by a fellowship of the Alexander von Humboldt foundation.

## AUTHOR CONTRIBUTIONS

A.K.C., M.J.C.G., Z.K., and E.V. performed experimental assays. P.P. and A.S. provided the HPLMax medium composition and fabrication protocol and gave experimental advice. A.K.C. and M.J.C.G. prepared HPLMax. F.C.A. and S.O. contributed to histology analyses. A.C.E. performed all bioinformatic analyses. A.K.C., A.C.E., M.J.C.G., and T.G.P.G. performed statistical analyses. K.M.H. gave advice on metabolic aspects of the study and cell culture conditions. T.F. provided and analyzed an additional data set to cross-check for HKG candidates. A.K.C., A.C.E., and T.G.P.G. prepared the figures. A.K.C. and T.G.P.G. wrote the manuscript. A.K.C., M.J.C.G., and T.G.P.G. designed experimental assays. T.G.P.G. conceived and supervised the study and provided the laboratory infrastructure and financial support. All authors read and approved the final manuscript.

## LEGENDS TO SUPPLEMENTARY TABLES

Supplementary Table S1: Composition and Fabrication of HPLMax

Supplementary Table S2: Pre-ranked fGSEA with C2, C5, and Hallmark gene sets from the MSigDB of DEGs in standard and refined cell culture conditions across 4 EwS cell lines

Supplementary Table S3: Comparison of DEGS being EWSR1::ETS positively or negatively correlated (|log2 FC|=1.5) in knockdown experiments of EWSR1::ETS in standard and refined cell culture conditions across 4 EwS cell lines

Supplementary Table S4: List of HKG candidates for gene expression comparison experiments in refined and standard EwS culture conditions

## METHODS

### Experimental models and subject details

#### Provenance of cell lines

The EwS cell line A-673 (RRID CVCL_0080) was obtained from American Type Culture Collection (ATCC, Manassas, USA), and MHH-ES-1 (CVCL_1411), SK-N-MC (CVCL_0530), and TC-71 (CVCL_2213) were obtained from the German Collection of Microorganisms and Cell cultures GmbH (DSMZ, Braunschweig, Germany). The EwS cell line TC-106 (CVCL_F531) was obtained from the Children’s Oncology Group (COG). Cell identity was regularly confirmed by single-nucleotide polymorphism (SNP) and/or short tandem repeat (STR) profiling. Genetic sex was ascertained from WGS data.

#### Culture conditions for cell lines

For standard culture conditions, all cell lines were cultured in RPMI 1640 medium with stable glutamine (Biochrom, Germany), supplemented with 10% FCS tested to be doxycycline (dox)-free (Sigma-Aldrich, Germany), and penicillin (100 U/mL) and streptomycin (100 μg/mL; Merck, Germany) in tissue culture flasks and plates (Sarstedt, Germany). For TC-106, culture dishes were coated with collagen coating solution (Sigma-Aldrich) and either directly used or stored at 4°C for max two weeks.

For refined culture conditions, all cell lines were cultured in HPLMax (protocol for fabrication see below; details of the composition are given in **Supplementary Table S1**), supplemented with 7% FCS tested to be dox-free (Sigma-Aldrich, Germany), and penicillin (100 U/mL) and streptomycin (100 μg/mL; Merck, Germany) in tissue culture flasks and plates (Sarstedt, Germany) coated with 2 mL of 1.5% high gel-strength agar (Serva) to generate ultra-low attachment conditions for spheroid formation^19^. Coated flasks were either directly used or stored at 4°C for max 2 weeks.

In both culture conditions, cells were incubated at 37°C and 5% CO2 in a fully humidified environment. Cells were subcultured in ratios 1:2 to 1:8 after detachment with trypsin/EDTA (Biochrom). Cultures were routinely tested for and found to be free of Mycoplasma contamination by nested PCR.

#### Spheroid seeding

For seeding of cells for spheroid formation, previously 2D grown cells in HPLMax supplemented with 7% FCS were used. Respective cell numbers of A-673/TR/shEF1 (1.2 × 106 cells), SK-N-MC/TR/shEF (1 × 106 cells), MHH-ES-1/TR/shEF1 (1.8 × 106 cells), TC-71/TR/shEF1 (1 × 106 cells) and TC-106/TR/shEERG (1.2 × 106 cells) were seeded in triplicates per condition (dox+/–) in five ml of HPLMax, 7% FCS, in agar-coated T-25 flasks (see description for refined culture conditions above). Cells were cultured for sphere formation for four days.

### Method Details

#### Fabrication HPLMax

For the refined cell culture conditions of EwS cell lines, we used a combination of two media which reflect metabolite concentrations of human plasma, namely HPLM and Plasmax^13,14^. Our combination, called HPLMax, is based on the readily available HPLM^13^ medium to which we added manually 13 additional metabolites, that are only included in the Plasmax recipe^14^. An overview on the compositions of HPLM and Plasmax^13,14^ is given in **Supplementary Table S1**. For the 13 additional metabolites, single stock solutions were prepared according to **Supplementary Table S1** and stored at –20°C. For preparation of HPLMax, the 13 Plasmax metabolite single stock solutions were combined, filter sterilized and added to HPLM^13^ as according to **Supplementary Table S1**.

#### Cell lines with EWSR1::ETS-knockdown

We used 5 previously generated EwS cell lines with inducible knockdown of the fusion oncogene^22^. Briefly, EwS cell lines were lentivirally transduced with an pLKO-Tet-On all-in-one vector containing a dox-inducible shRNA against the respective *EWSR1::ETS* fusion or against its constituting *ETS* parts (*FLI1* or *ERG*)^22^. Efficient gene silencing was confirmed by qRT-PCR and led to a remaining fusion gene expression of ∼20% and previously performed western blot confirmed knockdown of the respective EWSR1::ETS fusions at the protein level^22^. These established cell lines were regularly selected with 1 mg/mL puromycin (Invivogen). The shRNA mediated knockdown of the respective *EWSR1::ETS* fusion oncogenes was induced by addition of 1 μg/mL dox (Merck, Germany) to the cell culture medium. Dox was refreshed every 48 h. Afterwards, expression of the given *EWSR1::ETS* fusion and knockdown efficiency was assessed by qRT-PCR.

#### Proliferation assays

Proliferation of EwS cells was assessed by resazurin colorimetry^33^ or trypan blue exclusion^34^. In brief, a previously described drug screening by resazurin colorimetry in EwS^33^ was adapted to test cell growth in different FCS concentrations. A-673/TR/shEF1 (6 × 10^3^ cells), SK-N-MC/TR/shEF1 (5 × 10^3^ cells), MHH-ES-1/TR/shEF1 (5 × 10^3^ cells), TC- 71/TR/shEF1 (4 × 10^3^ cells) and TC-106/TR/shEERG (5 × 10^3^ cells) were seeded in 96-well plates (Sarstedt, Germany) in 1–10% FCS supplementation in RPMI, and read out was performed after 96 h incubation according to the previously published protocol^33^. Trypan-Blue exclusion was performed as described^34^. In brief, 3–20 × 10^4^ cells were seeded in 6-well plates in 10% or 7% FCS supplemented media and viable and dead cells were manually counted after 96 h.

#### Spheroid histology

Spheroids were seeded as described above. After 96 h of spheroid formation and additional 96 h of treatment with dox+/–, spheroids were harvested on day 9 after seeding. Spheroids were collected, incubated in 4% buffered paraformaldehyde (PFA) for 15 min at RT, then embedded into a cryomold (Sakura Finetek, USA) containing 3% agar. Once solidified, the agar pad was transferred into a histology cassette. The cassette was put in 4% buffered PFA overnight and transferred into a 50% alcohol solution the next day. Then, water extraction and embedding in paraffin was conducted by the DKFZ Light Microscopy Core Facility. Paraffin blocks were cut into slices of 3 μm, transferred onto microscope slides and stained with hematoxylin and eosin (H&E). For comparison, H&E stains of archival subcutaneous xenografts of wildtype TC-71 cells were used^9^.

#### RNA extraction and qRT-PCR

For RNA extraction, the NucleoSpin RNA kit from Macherey-Nagel (Duren, Germany) was used following the manufacturer’s protocol. Isolated RNA (1 mg, quantified with NanoDrop) was reverse transcribed applying the High-Capacity cDNA Reverse Transcription Kit (Thermo Fisher) following the manufacturer’s protocol. qRT-PCR reactions contained 9 μL 1:10 diluted cDNA, 10 μL SYBR Select Mastermix (Thermo Fisher) and 1 μL mix of forward and reverse primer (final concentration 0.5 mM) in a final volume of 20 μL. Reactions were carried out in duplicates or triplicates with a CFX Connect Real-Time PCR Detection System (Bio-Rad, Munich, Germany). Relative expression was calculated by the ΔΔCt method and normalized against either *ribosomal protein lateral stalk subunit P0* (*RPLP0)* or *RNA component of 7SK nuclear ribonucleoprotein* (*RN7SK)*. Stable expression of *RPLP0* in EwS cell culture conditions with RPMI medium has been shown in previously^22^ and our data suggested that *RN7SK* is among the suitable HKGs for comparison of gene expression levels in EwS cell lines in the refined culture conditions. The following primer sequences (eurofins Genomics, Ebersberg, Germany) were used: *RPLP0* forward: 5’- GAAACTCTGCATTCTCGCTTC -3’; *RPLP0* reverse: 5’- GGTGTAATCCGTCTCCACAG -3’; *EWSR1::FLI1* forward: 5’- GCCAAGCTCCAAGTCAATATAGC -3’; *EWSR1::FLI1* reverse: 5’- GAGGCCAGAATTCATGTTATTGC -3’; *EWSR1::ERG* forward: 5’- TCCAAGTCAATATAGCCAACAGAG -3’; *EWSR1::ERG* reverse: 5’- CTGTGGAAGGAGATGGTTGAG -3’; *RN7SK* forward: 5’- ATTGATCGCCACCCTTGAT -3’; *RN7SK* reverse: 5’- CTCTATCGGGGATGGTCGT –3’;

#### Transcriptome analyses

To study the EWSR1::ETS activity in refined culture conditions in EwS cells, microarray analysis was performed. Four selected EwS cell lines were seeded in triplicates for each condition (dox+/–) as described above for spheroid seeding and cultured for 8 days, with dox refreshed every 48 h. Spheroids were collected on day 9, pelleted, then resuspended in lysis buffer from NucleoSpin RNA kit (Macherey-Nagel,Germany). Further RNA extraction followed the manufacturer’s protocol. RNA (1 mg, quantified with NanoDrop) was checked for quality with a tape station using the RNA ScreenTape Analysis kit (Agilent) according to manufacturer’s protocol. RNA integrity number was ≥8.5 for all samples. Knockdown efficiency was assessed as described above and remaining expression of *EWSR1::ETS* was confirmed to be <30% of untreated samples. Samples were hybridized to Human Affymetrix Clariom D microarrays. The retrieved raw expression data were subjected to joint processing and analysis, including background correction, normalization, and gene-level summarization, with the previously generated data from standard culture condition^22^. Of note, experimental procedures of dox treatment, RNA extraction, and RNA concentration for microarray profiling were identical in both experiments conducted under refined and standard culture conditions to minimize impact of potential batch effects. Data were quantile normalized and summarized with Transcriptome Analysis Console (v4.0; Thermo Fisher Scientific) using the SST-RMA algorithm^35^. The data were annotated with the Affymetrix library for Clariom D array (version 2, homo sapiens) at gene level. This resulted in a list of 21,324 unique annotated genes. Exploratory data analysis was conducted with PCA to identify captured variation in the transcriptomic data. For identification of DEGs for each comparison of interest with significant and consistent fold changes (FCs) across all cell lines, normalized log2 gene expression values were subjected to DEG analysis with limma (v 3.58.1)^36^. DEGs were defined as having adjusted *P*<0.05 and arbitrarily |log2 FC| >1.5.

#### Pathway enrichment analysis

To identify enriched gene sets, all genes were ranked by their mean log2 FC for each comparison of interest and used as input for pre-ranked GSEA^37^. The analysis was conducted using fGSEA (v 1.28.0)^38^ with gene sets from the Molecular Signatures Database (MSigDB, h.all.v7.0.symbols.gmt, c2.cgp.v7.0.symbols.gmt, c5.all.v7.0.symbols.gmt). Enrichment plots for individual gene signatures were generated using the enrichplot package (v 1.22.0), clusterProfiler (v 4.10.1)^39^. Enriched gene ontology (GO) terms, defined as adjusted *P*<0.05 and normalized enrichment score |NES|>2 (10,000 permutations), were employed for network construction though the use of Weighted Correlation Network Analysis (WGCNA, v 1.72-5)^40^. In a first step, a binary matrix of GO terms and genes (in which 1 signifies that the gene is present in the GO term and 0 signifies that it is not) was built. Subsequently, for all possible pairs the Jaccard’s distance was calculated to generate a symmetric GO adjacent matrix. The dynamicTreeCut algorithm was employed to identify clusters of similar GO terms and the top 20% highest edges were selected for plotting. The highest scoring node within each cluster was designated as the cluster label (rName). The resulting nodes and network files were imported into Cytoscape (v 3.8.0) for network organization and visualization, as previously described^41^. For overrepresentation analysis (ORA) of GO terms DEGs, PANTHER was employed as classification system^42^.

#### Identification of HKG candidates in refined and standard EwS culture conditions

To identify suitable HKG candidates with a stable expression across culture techniques and EWSR1::ETS expression levels, the approach described by de Jonge *et al*.^26^ was adapted as follows: All genes retrieved after joint background correction, normalization, summarization, and annotation (as described above) were filtered for a maximum |log2FC| <2, with the maximum FC defined as the maximum minus the minimal gene expression value across all samples for each gene. Subsequently, all filtered genes were ranked according to their coefficient of variation (CV = standard deviation/mean). The genes with the smallest CV were considered top HKG candidates (see also **Supplementary Table S4**).

#### Statistical analysis and software

Statistical data analysis was performed using PRISM 10 software (GraphPad Software Inc., Ca, USA) or R (version 4.3.0) on the raw data. Two groups in functional *in vitro* experiments were compared with a two-sided Mann-Whitney test, if not indicated otherwise in the figure legends. Data are displayed as dot plots with vertical bars representing means and whiskers representing the standard error of the mean (SEM), if not indicated otherwise in the figure legends. Sample size for all *in vitro* experiments were chosen empirically. For generation of size proportional Venn diagrams, the DeepVenn web application was used^43^.

#### Data availability

Original Affymetrix transcriptome profiling data have been deposited at the Gene Expression Omnibus (GEO) under the accession code GSE270118. In addition, Affymetrix microarray data from our published Ewing Sarcoma Cell Line Atlas (ESCLA)^22^ was used for comparison (accession code: GSE176190).

---

**Supplementary Figure S1.**
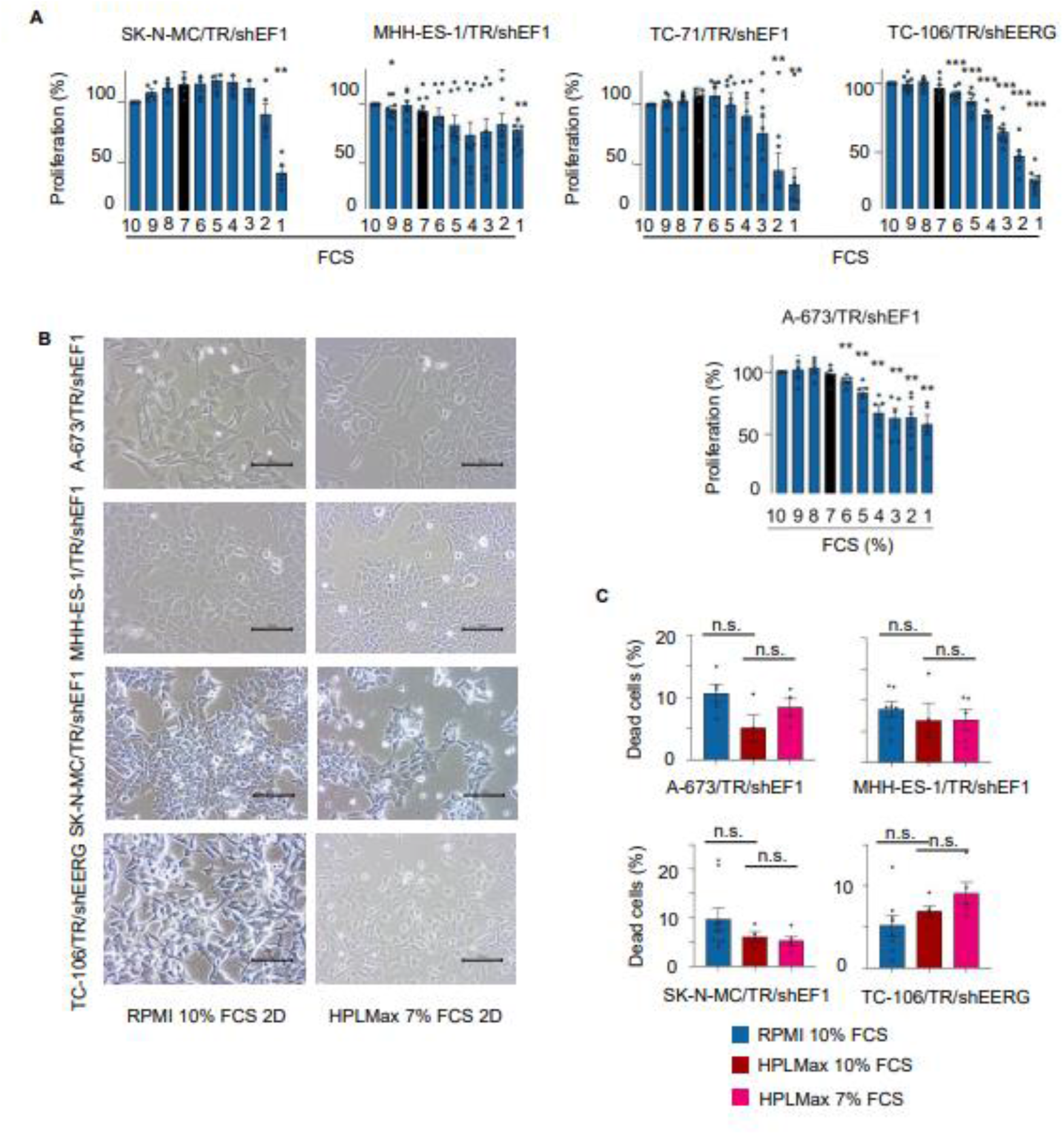
: Establishing physiologically optimized cell culture conditions for EwS cell lines **A** Proliferation assessment by resazurin cell viability assays in EwS cell lines in RPMI with FCS supplementation from 10% to 1%. Values were normalized to the proliferation in 10% FCS. Each dot represents a single biologically independent experiment. Vertical bars represent means and whiskers the SEM. Two-sided Mann-Whitney test. \**P*<0.05, \*\**P*<0.01, \*\*\**P*<0.001, black bars represent measurements at 7% FCS supplementation which was chosen as standard for further experiments in refined conditions. **B** Brightfield microscopy images of selected EwS cell lines grown in RPMI and 10% FCS (left) and in HPLMax and 7% FCS (right). Scale bars = 100 μm, 20× magnification. **C** Percentage of dead cells in comparison to viable cell count after 96 h of culture in EwS cell lines in RPMI, 10% FCS (left bar), HPLMax, 10% FCS (middle bar) and HPLMax, 7% FCS (right bar) by Trypan blue dye exclusion method. Each dot represents a single biologically independent experiment. Vertical bars represent means and whiskers the SEM. Two-sided unpaired student’s t-test. n.s.= not significant.

**Supplementary Figure S2:**
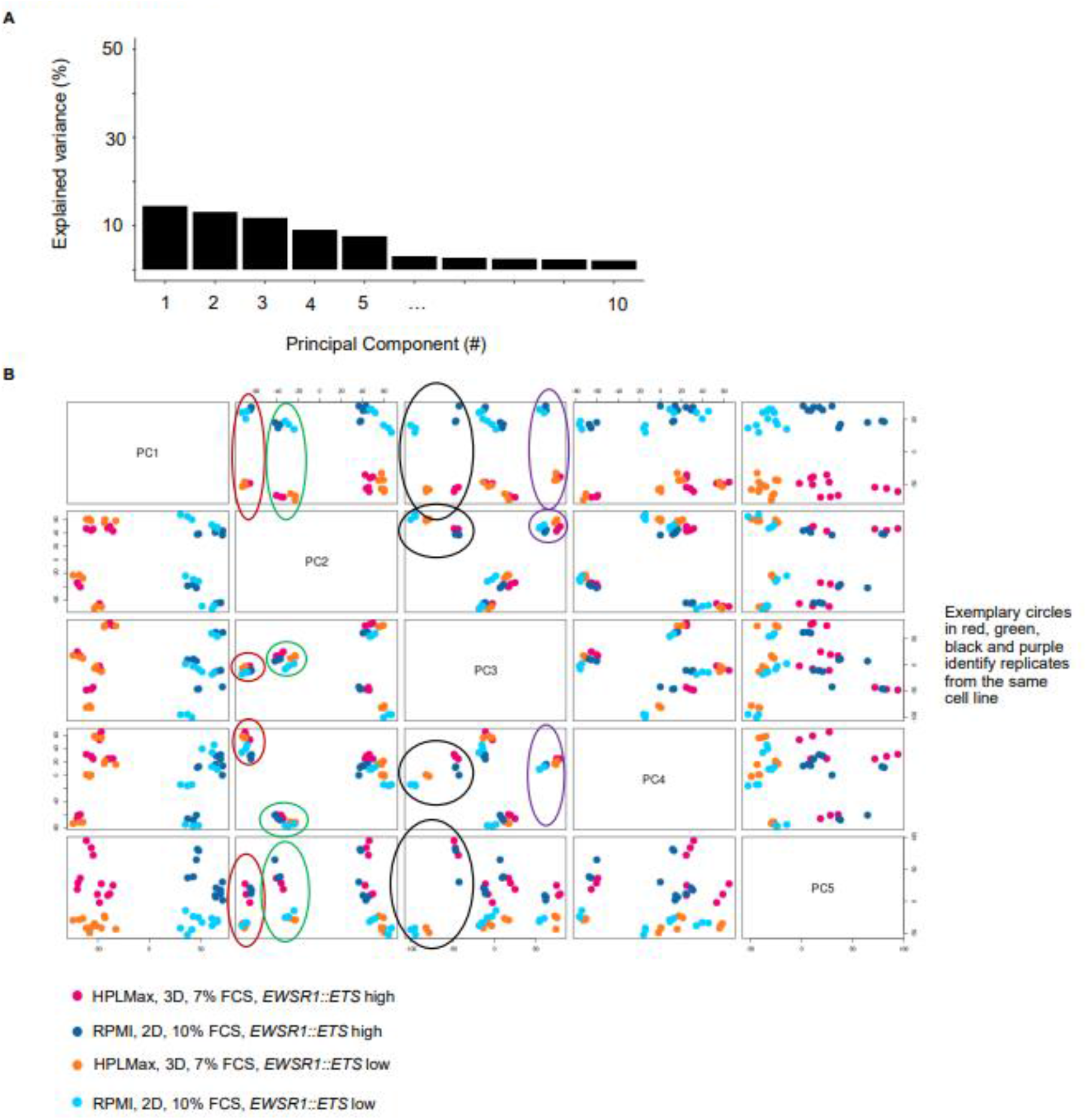
Physiologically refined EwS cell culture condition preserves cell line characteristics but alters transcriptional programs related to proliferation, differentiation, and metabolism **A** Scree plot of the PCA of the data from Figure 2A showing the explained variance (%) for top principal components (PC1-10) **B** Pairwise correlation plot based on PC1 to PC5. Each color represents one data set according to culture condition and *EWSR1::ETS* expression level. Exemplary circles in red, green, black, and purple identify replicates from the same cell line. **C** Enrichment plots from fGSEA of DEGs from Figure 2A. Padj = Bonferroni adjusted p value. C2 and Hallmark Gene Sets from the Human MSigDB Collections.

